# Broad antiviral effects of *Echinacea purpurea* against SARS-CoV-2 variants of concern and potential mechanism of action

**DOI:** 10.1101/2021.12.12.472255

**Authors:** Selvarani Vimalanathan, Mahmoud Shehata, Kannan Sadasivam, Serena Delbue, Maria Dolci, Elena Pariani, Sarah D’Alessandro, Stephan Pleschka

**Affiliations:** Pathology & Laboratory Medicine, University of British Columbia, Vancouver, Canada; Institute of Medical Virology, Justus Liebig University Giessen, 35392 Giessen, Germany; Center of Scientific Excellence for Influenza Viruses, National Research Centre, 12622 Giza, Egypt; Centre for High Computing, Central Leather Research Institute, Adyar, Chennai, India; Laboratory of Molecular Virology, Department of Biomedical, Surgical and Dental Sciences, University of Milano, 20133 Milano, Italy; Department of Biomedical Sciences for Health, University of Milano, 20133 Milano, Italy; Department of Pharmacological and Biomedical Sciences, University of Milano, 20133 Milano, Italy; German Center for Infection Research (DZIF), partner site Giessen

**Keywords:** Echinacea, SARS CoV-2, Variants of concern, Antiviral, Prevention, Spike protein, TMPRSS-2

## Abstract

**Background:** SARS-CoV-2 variants of concern (VOC) represent an alarming threat as they show altered biological behavior and may escape vaccination effectiveness. Some exhibit increased pathogenicity and transmissibility compared to the original wild type WUHAN (Hu-1). Broad-spectrum antivirals could complement and further enhance preventive benefits achieved through SARS-CoV-2 vaccination campaigns

**Methods:** The anti-coronavirus activity of *Echinacea purpurea* (Echinaforce® extract, EF) against (i) VOCs B1.1.7 (alpha), B.1.351.1 (beta), P.1 (gamma), B1.617.2 (delta), AV.1 (Scottish) and B1.525 (eta), (ii) SARS-CoV-2 spike (S) protein-pseudotyped viral particles and reference strain OC43 as well as (iii) wild-type SARS-CoV-2 (Hu-1) were analyzed. Molecular dynamics (MD) were applied to study interaction of Echinacea’s phytochemical markers with known pharmacological viral and host cell targets.

**Results:** EF extract broadly inhibited propagation of all investigated SARS-CoV-2 VOCs as well as entry of SARS-CoV-2 pseudoparticles at EC50’s ranging from 3.62 to 12.03 µg/ml. Preventive addition of 20 µg/ml EF to epithelial cells significantly reduced sequential infection with SARS-CoV-2 (Hu-1) as well as with the common human strain OC43. MD analyses showed constant binding affinities to Hu-1, B1.1.7, B.1.351, P.1 and B1.617.2-typic S protein variants for alkylamides, caftaric acidand feruoyl-tartaric acid in EF extract. They further indicated that the EF extract could possibly interact with TMPRSS-2, a serine protease required for virus endocytosis.

**Conclusions:** EF extract demonstrated stable antiviral activity across 6 tested VOCs, which is likely due to the constant affinity of the contained phytochemical marker substances to all spike variants. A possible interaction of EF with TMPRSS-2 partially would explain cell protective benefits of the extract by inhibition of endocytosis. EF may therefore offer a supportive addition to vaccination endeavors in the control of existing and future SARS-CoV-2 virus mutations.

## 1. Introduction

Replication of SARS-CoV-2 is known to be an inaccurate process because the viral polymerase introduces errors during replication of viral genome. This has led to the occurrence of novel variants, which differ from the WUHAN wild type strain (Hu-1), most vaccines are based on. Furthermore, recent findings revealed events of in-ter-lineage recombination of different SARS-SoV-2 variants co-infecting a single cell (Jackson, 2021, Banerjee, 2021). Since the genomic decryption of WUHAN-Hu-1 in January 2020 the evolution of new genetic lineages has been tracked globally and so far, thousands of different genomes have been collected by the National SARS-CoV-2 Strain Surveillance (NS3) system (Lu, 2020). Several new SARS-CoV-2 lineages are differing in survival fitness, infectivity, antigenicity and, most worryingly, in neutralization by vaccine-induced antibodies and sera (Zhou, 2021). The latter are carefully monitored as potential variants of concerns (VOCs) like the B1.1.7 (alpha-) variant, first detected in the United Kingdom bearing an N501Y mutation in the receptor binding domain (RBD) of the spike (S) protein. This variant rapidly spread from September 2020 on. It has meanwhile reached other regions of the world and shows reduced sensitivity to immune sera from Pfizer/BioNTech mRNA vaccinees and convalescents by factor 6 to 11, respectively (Collier, 2021, O’Toole, 2021). Notably, the novel strain exhibits a higher reproduction rate than pre-existing variants resulting in higher viral loads and longer infectious periods (Davies, 2021; Kidd and Kissler, 2021). Also, increased expression of inflammatory cytokines in nasal secretions was observed indicating pronounced pathology (Monal, 2021).

At the same time, other VOCs have emerged in South Africa (B1.351, gamma-variant) and in Brazil (P.1, beta-variant), both of which contain mutations at positions K417N/T and N501Y in the RBD, the domain that is essential for binding to human ACE-2 (hACE-2) receptors (Greaney, 2021; Castonguay, 2021). Again, a significantly reduced neutralization efficiency of sera from convalescent plasma against SARS-CoV-2 variants bearing the above-mentioned mutations was observed (Cele, 2021; Chen, 2021; Castonguay, 2021, 2021). Lately, in early 2021 the delta variant (B.1.617.2) emerged in Maharashtra, India, causing a massive pressure to the local healthcare and reached a prevalence of 87% by May 2021 (Salvatore, 2021). It shows increased lung cell entry and higher viral shedding/transmissibility in both vaccinated and unvaccinated individuals (Luo, 2021 Chen, 2021). As of July 2021, the delta lineage replaced most of previous variants, nowadays representing the predominant strain in USA and elsewhere producing a substantial number of vaccination breakthroughs (Christensen, 2021, Chia, 2021)

Originally, it was assumed that SARS-CoV-2 would have a minor propensity to mutate (an estimated 2 mutations per month), however as the pandemic lingers on, continuously new variants emerge, which potentially could escape effective immunization. The relevance of single point mutations for the cellular innate immune response might be lower than for the generation of the humoral immune response and epidemiological studies still reckon effective reduction of severe Covid-19 in fully vaccinated individuals (Cevik, 2021). At the same time, data draw a clearly less optimistic picture when it comes to mild to moderate diseases or delta variant transmission (Madhi,2021).

Thus, the search for robust preventive measures with low sensitivity towards genetic variations continues and might open up alternative strategies supplementing global vaccination endeavors. Especially the observed vaccine’s inefficiency in preventing asymptomatic and milder Covid-19 illnesses and virus transmission of the delta variant calls for additional solutions to reduce viral spreading since these cases account for the great majority of infections overall.

Antiviral activity has been identified in medicinal plants but the isolation of compounds/substances that exert the respective antiviral activity and manufacturing these into a product remains challenging (Vimalanathan, 2013). Plants produce a great variety of substances as secondary plant products, i.e. for their own defense against pathogens including viruses. Since many derivatives of a single chemical structure are produced naturally, the problem of point mutations in the virus genome and possible viral evasion is expected to be reduced (Pleschka, 2009). Likewise, antiviral activity against a wide series of common cold and highly pathogenic coronaviruses (CoV-229E, -MERS, SARS-1 and SARS-CoV-2) was previously demonstrated for an extract of *Echinacea purpurea* (Echinaforce® extract, EF) *in vitro* (Signer, 2020). Importantly, clinical effects on enveloped and endemic coronaviruses corroborated preclinical findings leading to recommendations on preventive use of Echinacea although modes-of-actions (MOA’s) are still poorly understood (Schapowal, 2020).

The current study aimed to investigate the antiviral potential of the *Echinacea purpurea* extract regarding the propagation of actual SARS-CoV-2 VOCs and to explore possible antiviral MOA’s. Virucidal activity against VOCs was studied at 3 different laboratories in parallel, using a standardized experimental protocol. We found strong and broad virucidal activity of EF extract against all investigated VOCs, including the predominant lineages alpha, beta, gamma and delta variant as well as the isolates eta and the Scottish one. The fact that (i) the entry of pseudotyped viral particles bearing SARS-CoV-2 S protein was inhibited at similar EF concentrations as the wild-type along with (ii) data from molecular modelling clearly point towards an interaction of EF extract with the viral SARS-CoV-2 S protein. Cellular pre-treatment with EF further reduced infectivity of SARS-CoV-2 virus and immunocytochemical staining and ELISA point towards possible inhibition of proteases (e.g. TMPRSS-2), crucial for viral cell entry. Overall, our results together with data from Signer (2020) indicate a broad antiviral activity against Coronaviruses and VOC’s highlighting multiple points of interference with viral infectivity by the EF extract and the contained known marker substances.

## 2. Materials and Methods

### 2.1 Test Material

In our experiments, we employed Echinaforce® (A.Vogel AG, Roggwil, Switzerland) a hydroethanolic extraction (65% v/v ethanol) of freshly harvested Echinacea purpurea produced according to Good Manufacturing Practices (GMP). Echinacea herb and roots are extracted at a drug-to-extract ratio, DER 1:11 and 1:12, respectively and combined at a final ratio of 95:5 (Batch no. 1053057). Ethanol was used for control matching the highest EF concentration in the individual experiment.

### 2.2 Cells and Viruses

African green monkey kidney epithelial cells clone C1008 (Vero-E6), ATCC (Cat# CRL-1586) and HEK293T cells, ATCC (Cat# CRL-3216) were maintained with growth media (DMEM supplemented with 10% fetal calf serum (FCS, Invitrogen) and 100 U/ml penicillin and 0.1 mg/ml streptomycin (P/S, Thermo). Human Nasal epithelial cells (HNEpC, Promocell Germany) and human bronchial epithelial cells (HBEpC, MatTek USA) were cultivated as per recommended protocol.

The following SARS-CoV-2 variant strains were used in this study: United Kingdom - B.1.1.7, South Africa - B.1.351, Brazil - P.1, India - B.1.617.2, Nigeria – B1.525, Scotland – AV.1 as well as the original SARS-CoV-2 (WUHAN-Hu-1) and endemic Coronavirus OC43.

### 2.3 SARS-CoV-2 pseudoparticle generation

Lentiviral pseudotyped viruses are generated in HEK293T cells which are transfected with a mixture of 0.6 µg p8.91 (HIV-1 gag-pol), 0.6 µg pCSFLW (lentivirus backbone expressing a firefly luciferase reporter gene), and 0.5 µg pcDNA3.1 SARS-CoV-2 Spike D614 in OptiMEM with 10 µl polyethyleneimine 1 µg/ml (Sigma) as previously described (Conceicao C, 2020). The viral supernatant was harvested at 48 and 72 h post transfection, centrifuged to remove cell debris, and stored at -80 °C. The viral pseudoparticles were concentrated by overlaying the clarified supernatant on 20% sucrose and centrifugation at 23,000 rpm for 2 h at 4 °C. The purified pseudoparticles were then aliquoted and stored at -80 °C.

### 2.4 Cell viability assay

HEK293T cells were seeded at a density of 2 × 10^4^ cells per well in 100 µl DMEM supplemented with 10% FCS in a 96-well plate. Following 48 h of incubation at 37 °C, 2-fold serial dilutions of EF at a starting concentration of 100 µg/ml were added to the cell monolayer. Cells were examined microscopically for any sign of compound-induced cytotoxicity after 48 h. The cell viability was also determined via incubation with CellTiter-Glo® (Promega) to measure the metabolic activity of inoculated and uninoculated cells for 1 h, and the luminescence was recorded after 10 min using GloMax® Discover Microplate Reader (Promega). For Vero E6 cells, toxicity assay has been carried out and reported earlier (Singer et al, 2020)

### 2.5 Virucidal Activity

Virucidal tests were carried out by the following facilities, that investigated the respective VOCs: B.1.1.7 (alpha), B1.351 (beta) and P.1 (gamma), Pleschka, University Giessen, Germany), B.1.1.7 (alpha), B1.617.2 (delta), AV.1 (Scotland) and B1.525 (eta) (Pariani, University Milano, Italy) OC43 (Vimalanathan, University British Columbia, Canada). SARS-CoV-2 pseudotype assays were performed by Viral Glycoproteins group, the Pirbright Institute, United Kingdom. All facilities used the following experimental approach to ensure comparability of results with gradual modifications. Alternatively, plaque reduction and cytopathic endpoint dilutions were used as indicated.

SARS-CoV-2 mutant suspensions were prepared at 250 plaque-forming units (pfu) per 80 µl of incomplete DMEM medium (DMEM without FBS). 80 µl of test substance was added at concentrations of 50, 25, 10, 5, 2 and 0.1 µg/ml (dry mass) or 0.2% EtOH as control (ethanol concentration of the highest EF concentration) and incubated for 1 h at room temperature (RT).

### 2.6 Plaque reduction assay

In 6-well tissue culture plates, Vero-E6 cells were subculture with DMEM growth media and incubated overnight at 37 °C with 5% CO2. On the next day, 250 PFU/well in DMEM of the different SARS-CoV-2 strains were incubated with different non-toxic concentrations of the EF plant extract for 60 min at RT. Afterwards, the Vero-E6 cells monolayer was washed once with 1x PBS and incubated with the different inoculum (virus/plant extracts mix) for 1 h at 37 °C, 5% CO2. Then the inoculum was removed and cells were incubated with Avicel media (MEM supplemented with P/S, 2% FCS, 1.25% Avicel (RC-591, Dupont)) for 48 or 72 h at 37 °C, 5% CO2. The Avicel media was subsequently removed and cells were washed 3x with PBS, fixed with 10% formalin for 60 min and stained (0.5% crystal violet, 20% methanol in dH_2_O) for 15 min at RT. To determine the 50% effective concentration (EC50), the plaque titers were calculated in percentage (% titer = (100/titer of untreated sample) x titer of extract/SC-treated sample) with the control (untreated) set as 100 % using GraphPad Prism version 9 (GraphPad Software, San Diego, USA). The EC50 is defined as the compound concentration that inhibited virus-induced plaque by 50%.

For the cytopathic effect (CPE) end-point method Vero-E6 cells were grown in 96 well plates as described above to reached 95 to 100% confluency. Virus suspension was prepared at 100 PFU/100 µl in incomplete media and added to cells at the indicated dilutions prior to incubation at RT for 1h. Growth media was removed and samples were transferred from 96-well plates to cell culture plates, returned to CO_2_ incubator then incubated for ±72 hours until CPE was examined visually.

### 2.7 Pretreatment of human airway epithelial cells

Primary human nasal epithelial cells (HNEpC) and primary human bronchial epithelial cells (HBEpC) were purchased from PromoCell GmbH, Germany, and MatTek Inc USA respectively. HNEpC and HBEpC cells were seeded (1×10^4^/well) in 12-well plates and cultured in growth media free of antibiotics until 70% confluency was attained. Cells were pretreated with increasing concentrations of EF (5 to 80µg/mL), untreated wells received media alone or media with 0.2% ethanol for 24, 48 and 78 h. At each time point, supernatant was removed, rinsed with PBS and infected with OC43 or SARSCoV-2 at the MOI of 1 and incubated for 72 h at 34 °C and 37 °C respectively in 5% CO_2_, CPE was monitored microscopically. For infectious viral titer analysis, at 72 h post infection (hpi), cells were scraped from each well and collected with supernatant and viral titer was determined on HCT or Vero-E6 cells. Briefly, one day prior to infection, HCT or Vero-E6 cells were seeded in 96-well plates at 1×10^4^ per well. Samples were serially diluted 2-fold with specific culture media containing 2% FBS and transferred to HCT or Vero-E6 cells, in 6 parallel wells (100 µL/well) and incubated for 72 h, CPE was observed, recorded and TCID 50/ml was calculated for each sample. The cytotoxicity of EF was tested in parallel on uninfected cells using the MTS assay (Promega).

### 2.8 Immunofluorescence staining

HNEpC cells were cultured on coverslips and pretreated as indicated in section (2.7). Pretreated human nasal epithelial cells were gently washed with cold PBS 3 times and fixed with 10% formalin for 10 min at RT. Fixative was then removed and cells were washed with PBS 3 times and permeabilized with 0.1% saponin in PBS for 5 minutes, permeabilization buffer was removed and cells were washed 3 times with PBST for 5 minutes each. Nonspecific antigens were blocked with 1% BSA in PBST for 1 h at RT. Then cells were incubated at 4 °C overnight with primary antibody (TMPRSS-2, Novus Biologicals cat# NBP1–20984) diluted to 5µg/ml in blocking buffer. Next day, primary antibody was aspirated and cells were washed 3 times with PBST, 5 min each, incubated with secondary antibody (Abcam, Ab150141) diluted to 1:250 in blocking buffer for 1 h at RT. Then cells were washed with PBST and mounted with 10 µl of aqueous mounting medium, Fluoroshield DAPI (Abcam, Ab 104139) for nuclear staining. Images were captured on Ziess Axio observer Z1 inverted fluorescent microscope (Zeiss, Germany) The fluorescent intensity was calculated using Imagej software (open source).

### 2.9 Molecular Modeling and Molecular Docking studies

The proper genome sequence of pandemic SARS-CoV-2 virus was retrieved from the National Centre for Biotechnology Information (NCBI) nucleotide database with the reference NC_045512.2. The available 3D crystal structures of various target SARS-CoV-2 proteins known as Papain-Like protease (PLpro), 3C-like protease (3CLpro), RNA-dependent RNA polymerase (RdRp), Spike (S) protein, human Angiotensin converting enzyme 2 (hACE2), Janus kinase 2 (JAK2) were obtained from protein data bank (PDB) (Berman, 2000) with the accession code 6W9C, 5R7Z, 7BV2, 6M0J_E, 6M0J_A and 2XA4 respectively. The optimum structures of other non-structural proteins namely NSP9, NSP13, NSP14, NSP15 and NSP16 were built suing homology modeling with appropriate templates using Swiss model (Waterhouse, 2018) and I-TASSER web-servers (Yang, 2015). The most active regions of the proteins were figured out using COACH meta-server (Yang, 2013) and outcomes were validated with CASTp web server (Tian, 2018). The antiviral and anti-inflammatory activities of compounds extracted from *Echinacea purpurea* against SARS-CoV-2 virus were examined by Molecular Docking studies using Schrodinger suite (Maestro) (Friesner, 2006). All the target proteins were processed by protein preparation wizard software (Schrodinger San Diago, Ca) and the ligands were constructed using LigPrep (Schrodinger San Diago, Ca). The OPLS3 force field with Extra Precision mode (XP) was performed to find the best position of ligands that adapt well in the cavity of the proteins.

### 2.10 Statistics

The experiments were conducted in triplicates and the given data are shown ± SEM or SD. One-way ANOVA followed by Tukey multiple comparisons test was performed using GraphPad Prism version 9 (GraphPad Software, San Diego, USA). The EC50 was determined using logarithmic interpolation.

### 2.11 Biosafety

All experiments performed with infectious VOCs were performed according to regulations for the propagation of BSL-3 viruses in a biosafety level 3 (BSL3) containment laboratory approved for such use by the respective local authorities.

## 3. Results

### 3.1 Pretreatment of VOCs prevents viral propagation at low, non-toxic EF concentrations

In a first approach to determine the antiviral potency of EF against the VOCs B1.1.7 (alpha), B.1.351.1 (beta), P.1 (gamma), B1.617.2 (delta), AV.1 (Scottish) and B1.525 (eta), variants were pretreated for 1 h with different, non-toxic concentrations (Singer, 2020) of EF (0 to 50 µg/ml) and subsequently this inoculate was used to infect Vero E6 cells (250 PFU/well). Viral propagation was then assayed via plaque assay at 48 or 72 h post infection (p.i.). EF exhibited a very potent and broad antiviral effect for all investigated SARS-CoV-2 VOCs (Figure 1). Complete prevention of viral propagation was observed at EF concentrations equal and higher than 25 µg/ml for all investigated VOC’s and across all laboratories (Figure 1). For all four laboratories, highly constant inhibitory concentrations were found with EC50 ranging from 5.37 to 12.03 µg/ml. One lab detected complete inhibition of P.1, B.1.351 and B.1.1.7 propagation at the lowest EF concentration of 1 µg/ml, which might be due to the fact that the virus stocks were derived from different sources for each lab. Results between labs were qualitatively highly consistent and show a strong inhibitory potential of EF extract irrespective of the different viral mutations.

**Figure 1.**
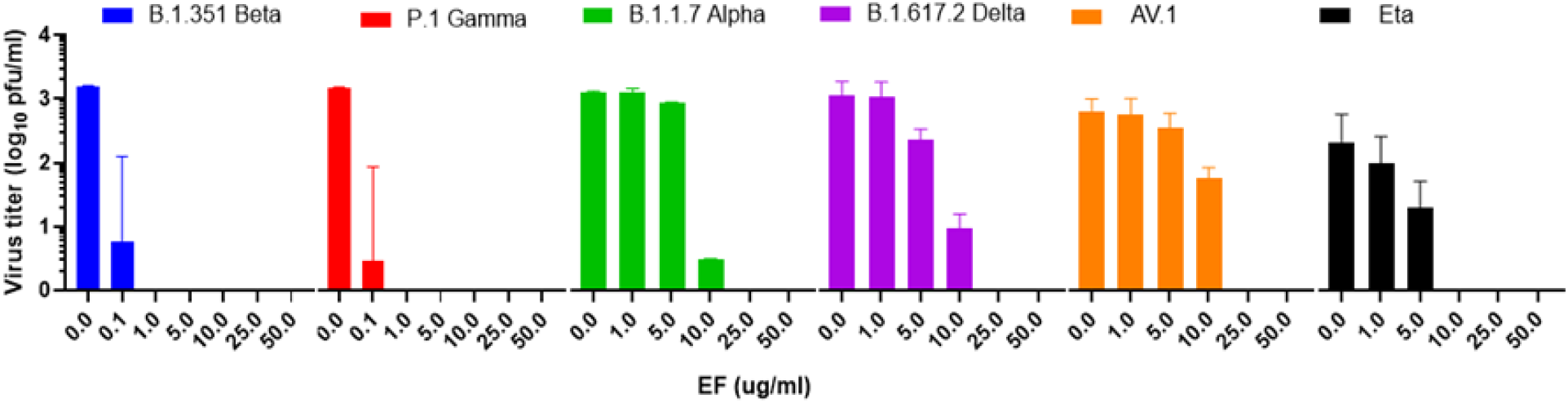
EF extract fully inhibited infectivity of all investigated variants of concerns (VOC’s) at concentrations ≥ 25 µg

### 3.2 Pretreatment of pseudoviruses expressing wild-type S protein prevents viral infection at low, non-toxic concentrations

SARS-CoV-2 pseudoparticles were generated via lentiviral-based pseudotyping approach, as previously described (Conceicao, 2020). EF showed a dose-dependent inhibitory activity against SARS-CoV-2 S protein-expressing pseudoparticles with an EC50 of 3.62 +/- 2.09 µg/ml (Figure 2). The fact that EF impaired the infectivity of both, SARS-CoV-2 and SARS-CoV-2 pseudoparticles at comparable concentrations highlights the S protein as a potential target for EF activity. Interestingly, the infectivity of SARS-CoV pseudoparticles was also inhibited by EF but at an EC50 of 16.7 +/- 10.15 µg/ml indicating a more potent activity towards SARS-CoV-2 over SARS-CoV.

**Figure 2.**
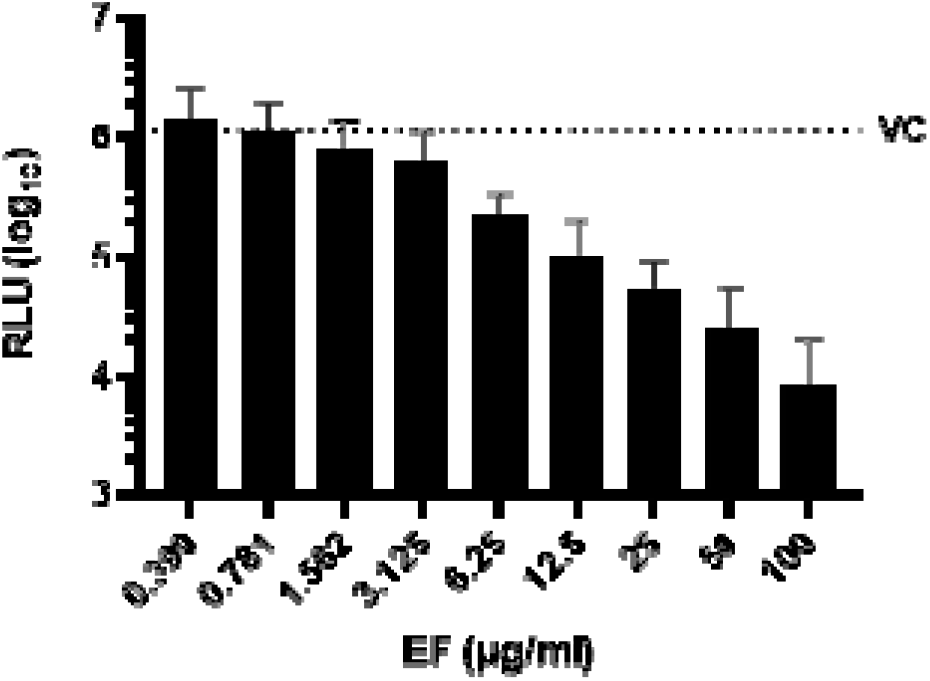
Inhibition of SARS-CoV-2 pseudoparticle infectivity by EF extract. Serial dilutions of EF extract were incubated with SARS-CoV-2 pseudoparticles for 1 h showing a dose-dependent inhibition of SARS-CoV-2 infectivity as measured by intracellular luciferase expression in mean relative light units (RLU). VC represents the RLU value of SARS-CoV-2 pseudoparticles-infected untreated cells. Data represent mean values ± standard deviations from three independent experiments performed in quadruplicates per experiment.

### 3.3 Pretreatment of primary nasal and bronchial epithelial cells impairs infection with OC-43 and SARS-CoV-2

Next to the preincubation of the viruses, primary human nasal epithelial cells (HNEpC) and primary human bronchial epithelial cells were preincubated with EF extract simulating a preventive treatment situation. EF at concentrations of 20 µg/ml completely reduced viral infections by SARS-CoV-2 (Hu-1) in both cell types as determined by CPE (Figure 3) end point assay after three days p.i.. Interestingly, in nasal epithelial cells infected with SARS CoV-2 CPE was detected only 5 days p.i., conversely, bronchial epithelial cells clearly showed CPE two days p.i..

**Figure 3.**
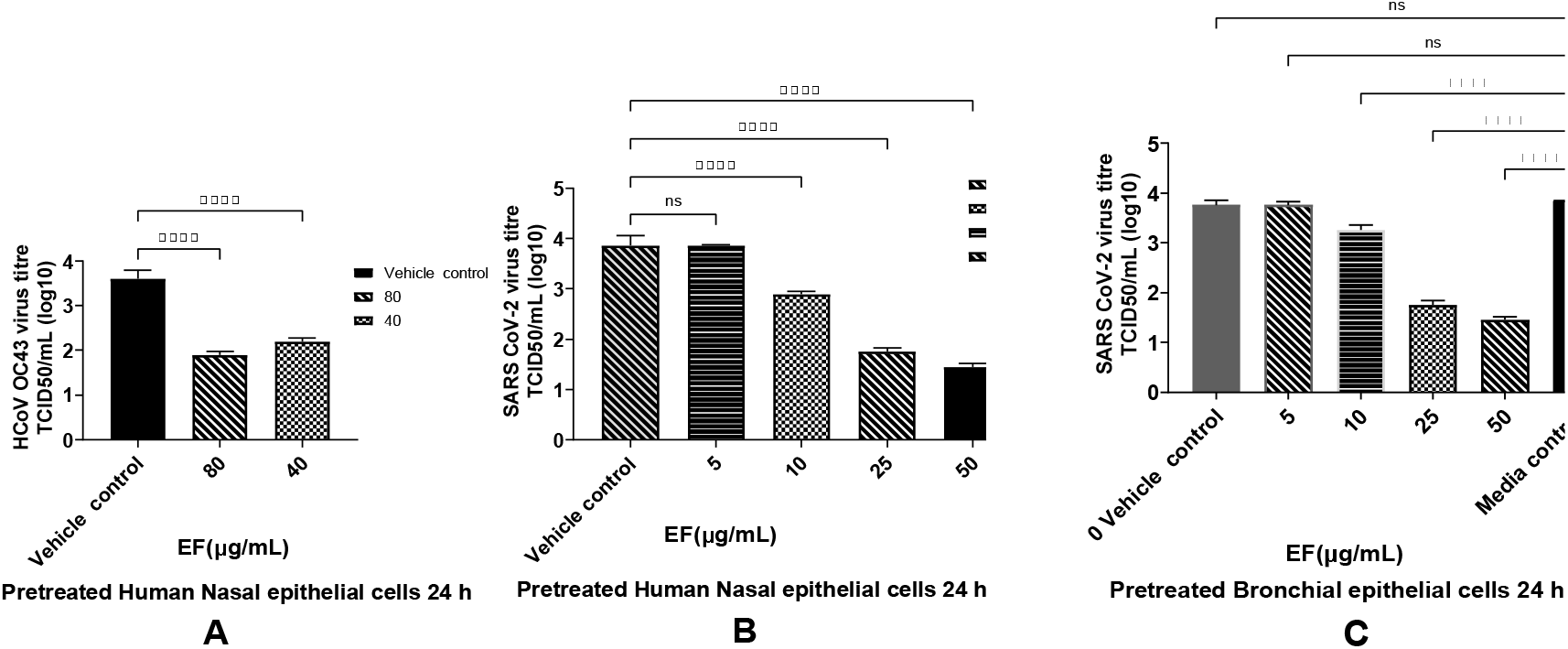
Infection preventive effect of EF on pretreated human airway epithelial cells. EF prevented OC43 (A) and SARSCoV-2 (B and C) infection dose dependently in EF pretreated HNEpC and HBEpC. Cells were pretreated with EF for 24 h, infected with virus at MOI of 1. Viral titer was measured at 72 h p.i.. Data result from 3 independent experiments. One-way ANOVA with Bonferroni multiple comparison correction was performed (****: p=<0.0001).

### 3.4 Molecular Docking (MD)

Herbal medicinal products typically contain a variety of pharmacologically active substances and in a first approach, several potential active pharmacological substances were investigated for their interaction with well-known viral and cellular proteins involved in SARS-CoV-2 infection. Therefore, various alkylamide derivatives, caftaric acid and 2-0-feruoyl-tartaric acid (totally 17 compounds) were tested for their docking potential against 12 different target proteins of SARS-CoV-2 virus (3CLpro, PLpro, RdRp, S-protein, NSP9, NSP13-16) and human cell proteins (ACE2, TMPRSS-2 and JAK2). The structure of the S protein and its receptor-binding domain (RBD, shown in red color) is represented in Figure 4. The RBD of the S protein contains i.e. 18 amino acid residues (K417, G446, Y449, Y453, L455, F456, A475, F486, N487, Y489, Q493, S494, G496, Q498, T500, N501, G502, Y505). Looking at the surface of the S protein, the active region of the protein has a cave-like domain, allowing ligands to adapt and fit into this domain (as shown in Figure 4).

**Figure 4.**
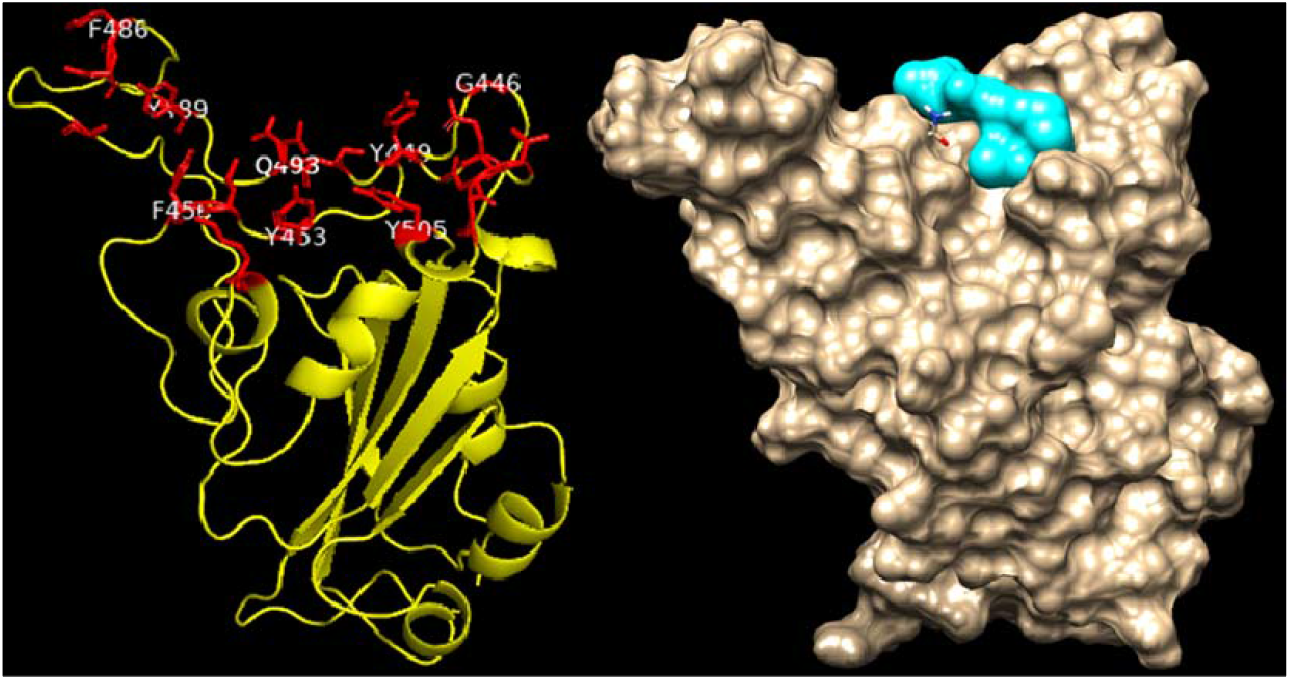
Structure of Spike protein of SARS-CoV-2 virus. The receptor binding domain (RBD) of the Spike protein is indicated in red color stick/blue.

The binding affinities of all 17 compounds towards 12 different target proteins of SARS-CoV-2 virus are shown (see Appendix A). For the compound dodeca-2E,4E,8Z,10E-tetraensaure-isobutylamid belonging to the alkylamide group, a higher binding energy of -7.6 kcal/mol with the S protein was identified. This is depicted in Figure 5, which shows hydrogen bonding interaction with the residues Gln 493 and Ser 494, Pi-Alkyl interaction with (Tyr 453, Tyr 495 and Tyr 505) and van der Waals interaction with residues Arg 403, Tyr 449, Gly 496 and Asn 501.

**Figure 5.**
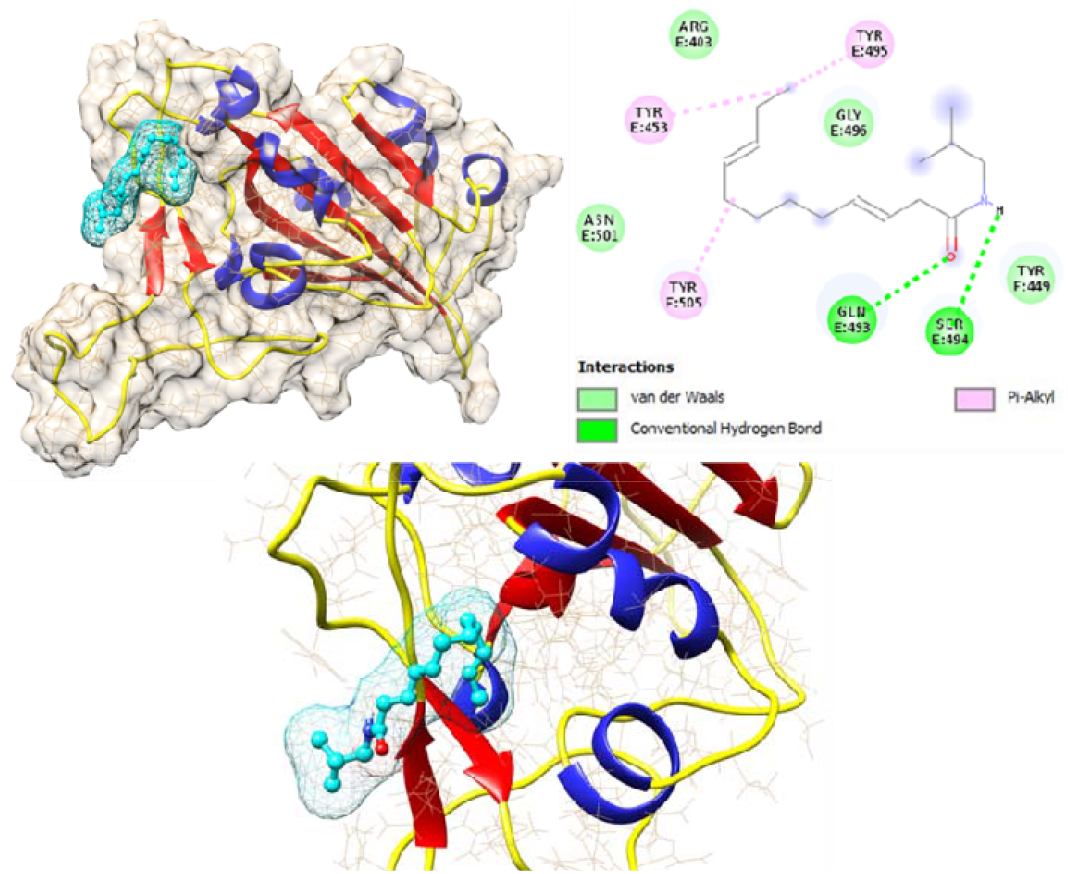
Interaction of dodeca-2E,4E,8Z,10E-tetraensaure-isobutylamid with wild type Spike protein of SARS-CoV-2 protein.

Furthermore, MD analysis also demonstrated that Alkylamides can interact with the RBD of the S protein. Caftaric acid and 2-0-feruloly-tartaric acid were both shown to have good binding affinity to the S protein and also to NSP’s with 5.1 / 5.7 kcal/mol and 7.4 /7.6 kcal/mol, respectively.

### 3.5 Echinacea’s Interaction with Pharmaceutical Targets for CoV Infection Process

Next, we investigated to which extent the molecular dynamic results obtained are applicable to a biological system and measured the effect of EF on the S protein interaction with the cellular binding receptor ACE2. To this point, an S protein (RBD)-coated ELISA plate (SARS-CoV-2 NeutraLISA, EuroImmune, Germany), originally developed to measure neutralizing antibodies, was incubated with EF extract, which reduced binding of biotinylated ACE2 receptors by 21.05 +/- 0.52 % at concentrations of 50 µg/ml, p<0.05 compared to the negative control (Figure 6). At least a partial blockade of the RBD could be deduced albeit interactions with other S domains are expected to fully explain the inhibitory effects of EF on CoV infectivity.

**Figure 6.**
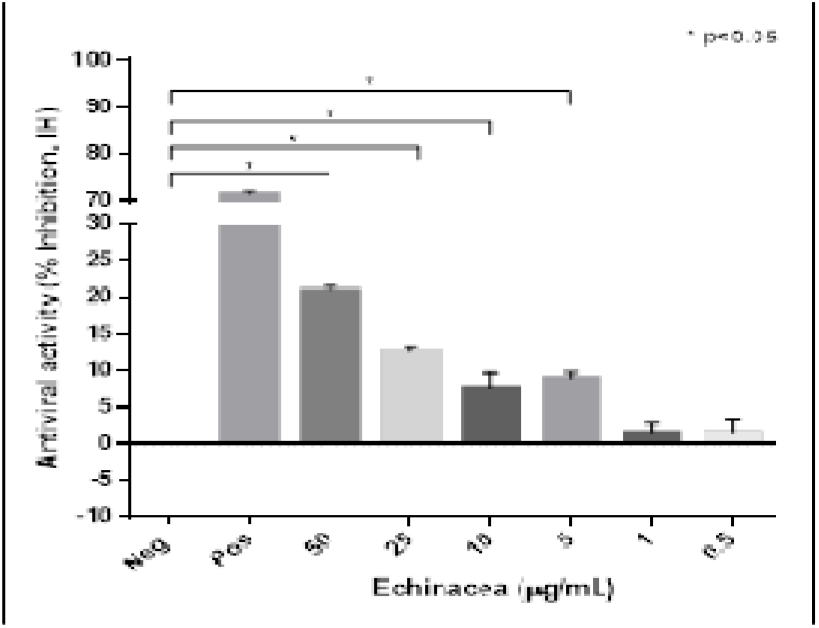
EF affects S protein binding to ACE2 receptor. S protein (RBD) coated ELISA plates were incubated with increasing EF concentrations and subsequently binding of biotinylated ACE2 receptors was quantified by measurement of chemiluminescence (% Inhibition, IH). As positive control ACE2 was co-incubated with immunized patient sera, which inhibited more than 71.5% of ACE2 receptor binding. EF at 50 µg/ml inhibited S protein binding, up to 21.0% to ACE2 (p<0.05)

Additionally, all compounds were tested as described in section 3.4 against x-ray structures of the S protein containing the mutations from the alpha, beta, gamma and delta VOCs and comparable binding energies as for wild type HU-1 were found. Data therefore indicate a very broad range binding activity of the different compounds to the S protein of different SARS-CoV-2 strains/VOCs, which is largely permissive for point mutations. Relevant interactions with S protein were further observed for other substances as well (e.g., caftaric acid), all of which appear to contribute to the overall bio-activity of EF.

### 3.6 Effect of EF on TMPRSS-2 expression

Furthermore, EF affected another cellular component. As shown by immunohistochemistry, treatment with EF extract at concentrations of 40 to 80 µg/ml resulted in significantly reduced expression of TMPRSS-2 in primary nasal epithelial cells (Figure 7 A), a result which was quantified by imagej software package (Figure 7 B). No effects of EF on the protease furin expression (needed for S protein activation) could be detected, even at EF concentrations above 160 µg/ml (data not shown). Taken together, the results obtained from the MD analysis, the ACE2/S protein binding analysis and the TMPRSS-2 expression study indicate a multifunctional MOA for EF, operating on various levels, i.e. partial interaction with the S protein and concomitant inhibition of cellular receptors required for viral attachment and invasion of the host, as well as impaired expression of an cellular proteas, which is essential for SARS-CoV-2 replication.

**Figure 7.**
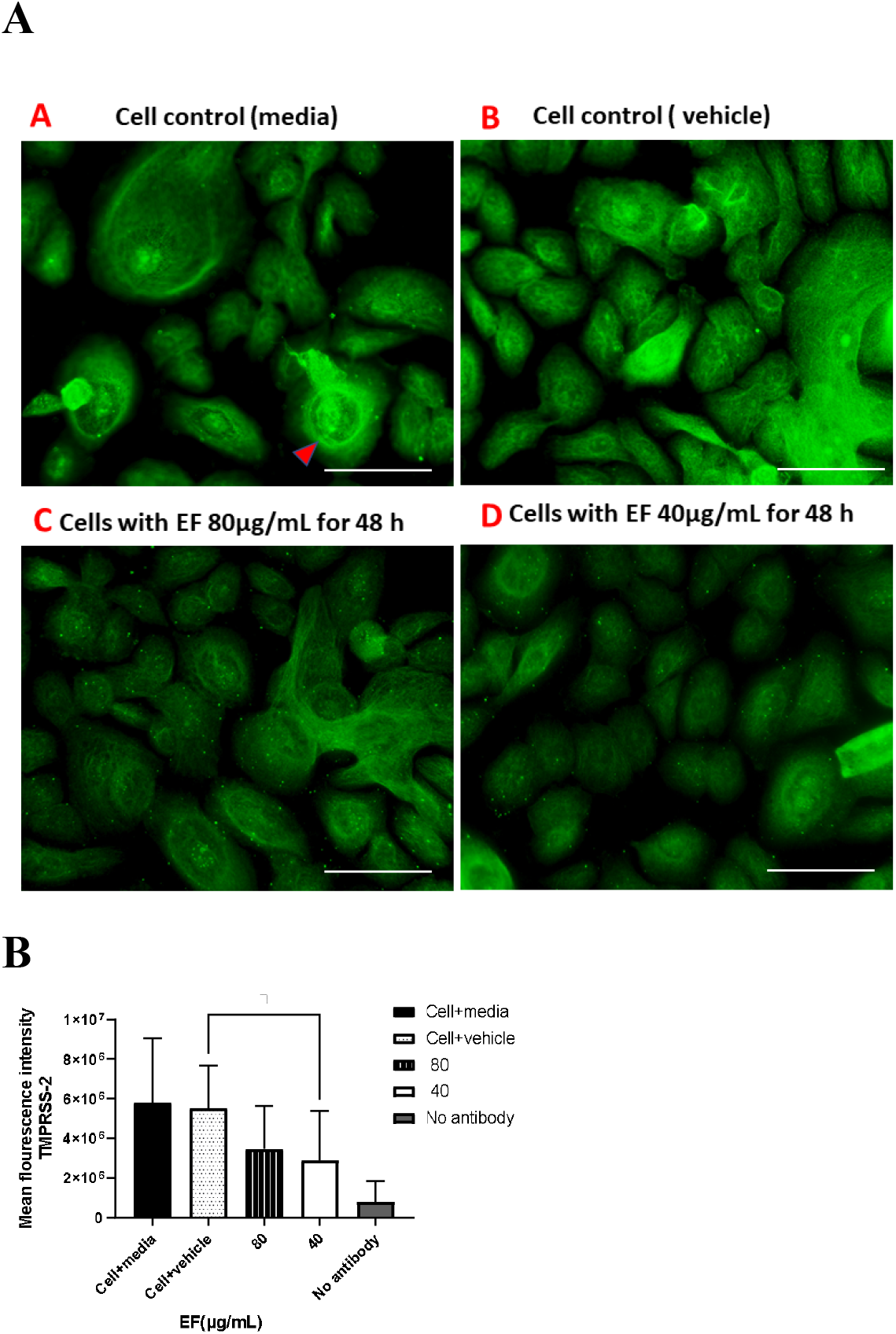
Inhibitory effect EF on TMPRSS-2 expression in EF pretreated HNEpC cells. Cultured HNEpC cells were treated with EF 80 and 40 ug/mL for 48 h. **(A)** Surface and intracellular immunofluorescent staining using an TMPRSS-2 antibody. **(B)** Quantification of the TMPRSS-2 signal. Graphs represent mean inflorescence values and ±SD of three independent experiments. Fluorescent intensity was calculated using imagej software package (open source). Scale bar =100µm.

## 4. Discussion

The Covid-19 pandemic has impressively demonstrated how viral pathogens can rapidly spill over from distant countries and spread world-wide within a few weeks’ time. Several vaccines using different immunization techniques were developed since and approximately one year after genomic characterization of SARS-CoV-2 in January 2020 first regulatory approvals were available. Nevertheless, another year elapsed until broad vaccination coverage e.g. of the Israelian population in order to achieve herd-immunity, was accomplished while most other countries have still not reached immunization rates of more than 50%, almost 2 years since occurrence of SARS-SoV-2. This delayed attainment of herd immunity gives the virus the possibility for further development through mutations and to evade containment by vaccines. In fact, viral variants and lineages develop faster than epidemiological studies are able to estimate real-life results for vaccination efficacy.

Thus, broadly effective and readily available antivirals are urgently needed, that are less susceptible to the spontaneous genetic variation frequently detected in viral respiratory pathogens not only in SARS-CoV-2. Multi compound extracts derived from medicinal plants might provide an option here as has been demonstrated for influenza viruses, known to quickly elicit resistance against oseltamivir treatment but not against treatment with *Echinacea purpurea* (Pleschka, 2009).

Our results together with earlier data from Signer (2020) infer a very broad virucidal activity of Echinaforce® extract against coronaviruses that comprises the actually circulating SARS-CoV-2 VOCs as well. Inhibition of infectivity exerted upon direct contact of virus with the extract but also a preventive treatment of cells provided a good level of protection against infection by the virus. Effective concentrations (EC50 < 12.03 µg/ml) varied marginally between different VOCs within the particular testing facilities. One laboratory generally measured antiviral effects at lower EF concentrations and MIC100 at 1 µg/ml for B.1.1.7, P.1 and B.1.351 virus, possibly reflecting the fact that the virus stocks were derived from different sources for each lab. Our results are in good agreement with previous findings for other enveloped viruses, where similar MIC100 were found for various influenza A virus strains (H3N2, H5N1, H7N7, H7N9 or influenza B virus) at EF concentrations below 50 µg/ml (Pleschka, 2009; Sharma, 2009).

Echinacea treatment of pseudoparticles specifically expressing SARS-CoV-2 spike D614 provided similar inhibitory effectiveness as with SARS-CoV-2 VOCs, suggesting an inhibitory action on the S protein, which may explain a MOA at the viral entry level as observed for influenza viruses (Pleschka, 2009). This hypothesis was further substantiated by ELISA experiments and molecular docking calculations revealing relevant binding affinities for a series of compounds in *Echinacea purpurea*, including alkylamides, its derivatives or caftaric acid. The fact that different substances show a certain level of binding not only to S protein but also to the serine protease (TMPRSS-2) or non-structural proteins (NSPs) indicates that the complexity of the whole extract might deliver the observed broad range inhibition in a concerted manner. Notably, different binding sites were identified for different compounds indicating synergistic effects of the complex substance mixture. As of yet, it appears impossible and meaningless to trace the antiviral activity down to a single substance in EF while still retaining its full spectrum benefits.

Remains the question regarding the clinical relevance of the presented findings, which, together with data from Signer (2020), infer a generic activity against coronaviruses overall. Evidence was gathered from three clinical studies, two on endemic CoVs and one comparative trial investigating SARS-CoV-2. For 4 months Jawad studied the effect of Echinaforce® on the incidence of respiratory tract infections in 755 volunteers. In this randomized, double blind, placebo-controlled, clinical study 54 viral infections occurred, of which 21 were caused by endemic coronaviruses (CoV-229E, CoV-HKU1 and HCoV-OC43) in the Echinacea group in contrast to 33 coronavirus infections in the placebo group (Jawad, 2012). Statistical significance was reached for the overall incidence of enveloped viruses (incl. coronaviruses, odds ratio OR=0.49; p=0.0114) rather than on the level of particular pathogens. Ogal and colleagues investigated the outcome of 4 months of preventive treatment with the same extract that we used and found a 98.5% decreased virus concentration in nasal secretions (p<0.05) in comparison with control treatment (Nicolussi, 2021). Again, significantly fewer enveloped virus infections (including coronaviruses) occurred with EF-treatment overall (OR=0.43; p=0.0038), substantiating the relevance of antiviral effects *in vivo*. Of note, in both studies participants kept the Echinacea formulation in the mouth for a few seconds prior to swallowing to enhance local antiviral effects. A very recent study was carried out during Covid-19 pandemic, which routinely collected naso/oropharyngeal samples during 5 months preventive treatment with EF extract or control. Here, Echinaforce® significantly reduced SARS-CoV-2 incidences from 14 to 5 infections (RR=0.42, p=0.046) and overall viral loads by more than 2.1 logs, corresponding with a >99% viral reduction (p=0.04). Finally, the time to become virus-free was significantly shortened.

The above presented data clearly indicate the clinical relevance of *in vitro* antiviral effects observed for *Echinacea purpurea* extract. This is highly important because all coronaviruses tested so far were inactivated by EF extract and the applicability of results to the *in vivo* situation would imply a general protective benefit not only against endemic strains, but also to newly occurring coronaviruses including their VOCs.

## 5. Conclusions

EF extract demonstrated broad and stable antiviral activity across 6 tested VOCs, which is likely due to the constant affinity of the contained phytochemical marker substances to all spike variants. A potential interaction of EF with TMPRSS-2 would partially explain cell protective benefits of the extract by inhibition of viral endocytosis. EF may therefore offer a supportive addition to vaccination endeavors in the control of existing and future virus mutations.

## Author Contributions

Conceptualization, S.V., M.S., D.S., P.E. and S.P.; methodology, validation formal analysis, resources, data curation, S.V., M.S.; D.S., K.S. P.E. software, S.V., M.S., K.S., D.S.; writing—original draft preparation, S.V., M.S., K.S. D.S. and S.P.; writing—review and editing, all authors.; visualization, S.V., D.S. K.S. and S.P.; supervision, S.V., K.S., P.E. and S.P.; funding acquisition, M.S., S.V., P.E. and S.P. All authors have read and agreed to the published version of the manuscript.

## Acknowledgments

We would like to appreciate Prof. Sandra Ciesek, Frankfurt, Germany, for kindly providing the SARS-CoV-2 variant strains South Africa - B.1.351 and Brazil - P.1, and Prof. Eva Friebertshäuser, Marburg, Germany, for providing United Kingdom variant strain - B.1.1.7. We kindly thank Dr. Dalan Bailey and Dr. Ahmed Mohamed from the Pirbright Institute, Ash Road, Pirbright, United Kingdom for carrying out studies on SARS-CoV-2 pseudotype virus expressing spike protein and for their contribution to the manuscript and critical review.

