## Supplemental Table 1 for "Broad antiviral effects of *Echinacea purpurea* against SARS-CoV-2 variants of concern and potential mechanism of action"

**Appendix 1.** The binding affinity of compounds of *Echinacea purpurea* against various target proteins of SARS-CoV-2 virus and of human cells in kcal/mol using Schrödinger XP Glide software package.

| Name of Echinacea purpurea Compounds | 3CL pro | PLp ro | Rd Rp | S-prote in | NS P9 | NSP 13 | NSP 14 | NSP 15 | NSP 16 | AC E2 | TMPR SS2 | JAK-Janus Kianse |
|---|---|---|---|---|---|---|---|---|---|---|---|---|
| Undeca-2E,4Z-dien-8,10-diinsaure-isobutylamid | -5.7 | -4.6 | -5 | -5.4 | -4.6 | -5.5 | -6.5 | -5.9 | -5.6 | -4.3 | -5 | -5.8 |
| Undeca-2Z,4E-dien-8,10-diinsaure-isobutylamid | -5.7 | -4.7 | -5 | -5.3 | -4.4 | -5.6 | -7 | -4.8 | -5.8 | -5.9 | -5.2 | -5.8 |
| Dodeca-2E,4Z-dien-8,10-diinsaure-isobutylamid | -5.1 | -5.6 | -5.2 | -5.2 | -4.5 | -5.8 | -7.4 | -5.8 | -6 | -5.6 | -5 | -6.1 |
| Undeca-2E,4Z-dien-8,10-diinsaure-2-methybutylamid | -5.3 | -5.3 | -5.1 | -5.2 | -4.5 | -5.7 | -6.7 | -5.6 | -6.1 | -5.9 | -5.5 | -5.8 |
| Dodeca-2E,4E,10E-trien-8-insaure-isobutylamid | -5.2 | -5.4 | -5.4 | -5.3 | -4.9 | -5.9 | -7.3 | -6.1 | -5.8 | -5.7 | -5.6 | -6.1 |
| Trideca-2E,7Z-dien-10,12-diinasaure-isobutylamid | -4.8 | -4.4 | -5.2 | -5.3 | -4.5 | -5.6 | -6.6 | -5.8 | -5.6 | -4.4 | -5.3 | -5.9 |
| Dodeca-2E,4Z-dien-8,10-diinasaure-2-methybutylamid | -5.2 | -5.4 | -5.2 | -6.9 | -4.1 | -6.4 | -7.1 | -5.5 | -5.8 | -6.3 | -5 | -6.3 |
| Dodeca-2E,4E,8Z,10E-tetraensaure-isobutylamid | -5.2 | -4.7 | -5.2 | -7.6 | -4.7 | -5.9 | -7.6 | -6.2 | -6.1 | -4.7 | -5.2 | -5.8 |
| Dodeca-2E,4E,8Z,10Z-tetraensaure-isobutylamid | -5 | -5.3 | -5.1 | -6.0 | -4.6 | -5.9 | -6.6 | -6.2 | -5.8 | -5.5 | -5.4 | -5.9 |
| Dodeca-2E,4E,8Z-triensaure-isobutylamid | -4.9 | -5.3 | -5.3 | -6.0 | -4.3 | -5.7 | -6.3 | -5.9 | -5.6 | -4.4 | -5.3 | -5.5 |
| Dodeca-2E,4E-diensaure-isobutylamid | -4.7 | -4.3 | -4.8 | -5.0 | -4.3 | -5.5 | -6.5 | -5.1 | -5.6 | -5.6 | -5.3 | -5.7 |
| 7-Hydroxy-Dodeca-2E,4E,8Z,10E-tetraensaure-isobutylamid | -5.4 | -4.8 | -5.7 | -4.8 | -4.5 | -6 | -7.1 | -6.5 | -6.2 | -4.8 | -5.8 | -5.9 |
| Undeca-2E,4Z-dien-8,10-diinsäure-isobutylamide | -5.2 | -5 | -5 | -5.6 | -4.8 | -5.7 | -6.9 | -5.9 | -5.7 | -5.6 | -5.3 | -5.8 |
| Pentadeca-2E,9Z-dien-12,14 diinsaure-isobutylamide and hydroxylated derivates | -5.1 | -4.3 | -4.6 | -4.9 | -4.3 | -5.8 | -6.8 | -5.2 | -5.4 | -4.8 | -4.9 | -5.7 |
| Trideca-2E,7Z-dien-10,12-diinsäure-isobutylamid | -4.9 | -5.3 | -4.8 | -5.5 | -4.6 | -6.3 | -7.3 | -5.6 | -5.5 | -6 | -5.5 | -6 |
| Caftaric acid | -6.8 | -6.4 | -6.6 | -5.1 | -5.1 | -7.1 | -7.4 | -6.6 | -7.3 | -6.4 | -6.6 | -6.8 |
| 2-0-feruloly-tartaric acid | -6.4 | -6.3 | -6.6 | -5.7 | -4.6 | -7.1 | -7.6 | -6.2 | -7.4 | -5.9 | -6.4 | -6.6 |
